# Remembering the opponent: neuronal activation associated with social memory in Zebrafish

**DOI:** 10.1101/2025.02.24.639917

**Authors:** Luciano Cavallino, María E Pedreira, Andrea G Pozzi, María F Scaia

## Abstract

Learning and remembering an opponent and the characteristics of a previous encounter may allow individuals to modify their behavior based on that acquired information. In the present study, using a long-term memory paradigm associated with social interactions, we aimed to evaluate the neuronal activation underlying individual recognition and memory of a previous agonistic encounter in male zebrafish. By quantifying a marker of neuronal activation, the immediate early gene c-fos, we compared the activation in different telencephalic nuclei belonging to the social decision-making network. Two dorsal nuclei, the medial (Dm) and the lateral (Dl), and two ventral nuclei, the ventral (Vv) and the dorsal (Vd), were evaluated by quantifying c-fos protein by immunohistochemical techniques. Two agonistic encounters between the same pair of opponents were performed, and the number of c-fos-immunopositive cells was quantified immediately after the second encounter in fish that had been treated with an amnesic agent after the first encounter (MK-801) and untreated individuals (Water), as well as a Control group with no physical interaction. We found that the Vv nucleus showed lower activation in individuals not exposed to physical interaction than those exposed (MK-801 and Water treatment). The Dl nucleus showed a higher activation only in individuals previously treated with MK-801, who would face an unremembered opponent, but not in those treated with water, which recognized the opponent. Our study provides evidence as a first step to understanding the neuronal processes underlying individual recognition and retrieval of an opponent.

**Summary statement:** Our study provides new information about the neuronal activation related to individual recognition and social interaction in zebrafish

## Introduction

Social experience can modulate behaviors in individuals and modify the interaction between conspecifics (Johnsson, 1997). Complex animal social behaviors involve different cognitive skills, such as observational learning, social learning, and individual recognition. In this sense, long-term memory formation could be beneficial in improving future responses adjusted to past experiences (Huntingford, 2013). During an aggressive interaction, individuals can sense multisensory cues, learn from their encounters, and remember different features of them. Consequently, animals can change their behaviors accordingly and prevent energy waste and injuries (Smith and Price 1973; Shettleworth, 2001). For this to occur, it is necessary to recognize and remember the opponent and recall this information in subsequent encounters. Social learning has been suggested to shape the evolution of social behavior (Fernald 2017), and considering that cognitive skills need to be transmitted and supported by neural systems for social systems to evolve, it is essential to characterize the neuronal activation and assess the brain nuclei involved in this kind of memory.

Associated with capabilities, we can mention two very important neural circuits in mammals, the mesolimbic reward system and the social behavior network, which evaluate stimulus salience and regulate social behaviors. Both constitute the Social Decision-Making Network (SDMN) (O’Connell & Hofmann, 2011). The social behavior network comprises the preoptic area, the anterior hypothalamus, the ventromedial hypothalamus, the periaqueductal gray, the medial amygdala, the lateral septum, and the bed nucleus of the stria terminalis (BNST). It has been observed that these regions are important in regulating aggressive, reproductive, and parental behaviors in mammals. Meanwhile, the mesolimbic reward system consists of the striatum, nucleus accumbens, the ventral pallium, the basolateral amygdala, the hippocampus, the ventral tegmental area, and shares the medial amygdala, lateral septum, and BNST with the social behavior network. It is suggested that this network is responsible for evaluating the salience of perceived stimuli to generate the appropriate behavioral response (O’Connell & Hofmann, 2011). In teleost fish, a consensus has emerged from neurophysiological, behavioral, anatomical, developmental, and molecular evidence suggesting that different areas in the telencephalon and hypothalamus are homologous to specific brain regions in mammals (Perathoner et al., 2016). Regarding ventral telencephalic areas, while the ventral (Vv) and lateral (Vl) zones are the suggested homologies for the mammalian lateral septum, the dorsal zone (Vd) is the homologous of the striatum and partly with the nucleus accumbens. When referring to the dorsal telencephalon, while the medial zone (Dm) in teleosts is the suggested homology for the ventral pallium in mammals (basolateral amygdala), the posterior zone (Dp) is the homologous of the lateral pallium, and the lateral zone (Dl) of the medial pallium (hippocampus). These homologies were later confirmed with evidence of functional connectivity, evaluating how certain regions responded to particular stimuli and the interconnection between the different regions and known cell populations. The study of gene expression in various brain areas in fish added more evidence to these homologies, comparing the expression of specific characteristic genes of different regions in mammals (O’Connell & Hofmann, 2011). Finally, activation and lesion experiments in various brain regions allowed a comparison of the effects in fish with the impact of activating or lesioning known areas in mammals, thus adding neurophysiological and behavioral evidence of the proposed homologies (Perathoner et al., 2016). In line with these results, many studies have assessed the activation of these telencephalic nuclei in the zebrafish *Danio rerio* through the quantification of various molecular markers of neuronal activation (Teles et al., 2015; Kareklas et al., 2023; Akinrinade et al., 2023). Other studies evaluated the activation of these nuclei associated with social interactions and aggression and the co-activation or co-repression between them (Scaia et al., 2022).

A widely used technique to evaluate neuronal activation is quantifying immediate early genes (IEGs). While the expression of these genes is not specific to neurons, it has been observed in neuronal cell cultures how the transcription of early genes was induced by electrical stimulation and by activation through neurotransmitter analogs, such as nicotine (Morgan and Curran, 1986; Bartel et al., 1989). The same effect was observed when the nervous system was stimulated *in vivo*, inducing the expression of c-fos, one of these types of genes (Mizuno et al., 1989). In this way, it has been observed that the expression of IEGs occurs rapidly and transiently following synaptic stimulation (Okuno, 2011). Specifically, the expression of several of these genes, including c-fos, was rapidly increased following neuronal activation, whether through pharmacological treatment or sensory stimulation (Minatohara et al., 2016). The quantification of c-fos has been widely used as a marker of neuronal activity in various species, including zebrafish (Wilson et al., 2002; Hudson, 2018; Chen et al., 2016). A commonly used technique is the measurement of the c-fos protein by immunohistochemical methods (Chatterjee et al., 2015). Chatterjee and colleagues observed that in zebrafish, stimulation through exposure to caffeine correlated with an increase in the measurement of the c-fos protein in various brain regions (Chatterjee et al., 2015) and Chaudhuri and colleagues evaluated the temporal patterns of c-fos protein expression in the rat neocortex, finding, through immunohistochemical techniques, that the expression of this protein could be observed between 30 minutes and 2 hours after the stimulus, with a peak at 90 minutes (Chaudhuri et al., 2000). Considering these previous pieces of evidence, many studies in zebrafish use time points of 60 minutes to observe peaks in protein expression, although it can also be observed at 30 minutes (Von Trotha et al., 2014; Chatterjee et al., 2015).

Learning processes and behavioral changes can persist over time through long-term memory formation, which requires synaptic plasticity changes dependent on molecular signaling cascades. These changes can strengthen specific synaptic connections and discrete brain networks (Asok et al., 2019). In this way, particular brain areas can be related to certain types of learning and memory by evaluating neuronal activation, the strengthening of neural circuits, and the effect of inhibiting or activating specific brain regions.

In previous experiments, we determined that adult male zebrafish resolved a second agonistic encounter with lower levels of aggression only when this second encounter was with the same opponent. If presented with a novel opponent, they resolved the second encounter with aggression levels similar to the first (Cavallino et al., 2020). These results were observed with intervals between encounters of 24 and 48 hours. Additionally, we observed that if, after the first encounter, the individuals were treated with an amnesic agent, MK-801, which is a non-competitive NMDA receptor antagonist, they did not decrease their aggression in the second one, behaving as if they were facing a novel opponent (Cavallino et al., 2024). The present study analyzed brain activation associated with individual recognition and long-term memory in zebrafish. We evaluated four telencephalic nuclei related to social interactions, recognition, and memory by quantifying c-fos protein expression. Activation and correlation among these nuclei were compared in individuals exposed to a subsequent encounter with a known and remembered opponent, exposed to a subsequent encounter with an unremembered opponent, and exposed chemically but not allowing the physical interaction. We found evidence of differential activation of these nuclei related to exposure or non-exposure to social interaction and the presence or absence of long-term memory associated with recognizing and remembering opponents.

## Material and method

### Ethics statement

The procedures used in this study followed the institutional guidelines for the use of animals in experimentation (Comisión Institucional para el Cuidado y Uso de Animales de Laboratorio, Facultad de Ciencias Exactas y Naturales, Universidad de Buenos Aires, Protocol 75b/2021) and were in accordance with National regulations (Comité Nacional de Ética en la Ciencia y la Tecnología). All procedures complied with the Guide for Care and Use of Laboratory Animals (eighth ed. 2011, National Academy Press, Washington, p. 220.) and the ARRIVE guidelines.

### Animal and maintenance

Adult zebrafish males, *Danio rerio* (Hamilton-Buchanan 1822), (n=21, weightL=L0.18±0.01 g, standard lengthL=L2.40±0.03 cm; mean ± SEM) were obtained from commercial aquaria in Buenos Aires, Argentina. One month before the experiments, individuals were acclimated in 20-L aquaria (densityL=L1 individual/L; 4 different aquariums): 25–26 °C, pHL=L7.5/7.8 and a photoperiod of 14 h light: 10 h dark cycle (Avdesh et al. 2012). They were fed twice daily with commercial food (Tetra®). For the present study, we analyzed the brains from a subset of samples from the experiments described in Cavallino et al., 2024.

### Experimental treatments

In the present study, we analyzed individuals from three experimental treatments (Fig.1):

#### Control

A pair of adult male zebrafish of similar size was placed in the experimental tank, separated by a barrier with visual but not chemical isolation. They remained with no physical interaction for 48 hours until they were euthanized by cold shock and decapitation, immediately fixing the heads as described in the following section.

#### Water Treatment

A pair of adult male zebrafish of similar size was placed in the experimental tank, separated by a barrier that isolated them visually but not chemically. After 24 hours of isolation, the barrier was removed, and their interaction was recorded for 30 minutes. After this time, they were separated and placed for 60 minutes in a treatment tank with pure water. Then, they returned to their respective experimental tank, where they remained isolated for 24 hours until the barrier was removed, allowing a new interaction between the same opponents. After the 30-minute encounter, they were immediately sacrificed by cold shock and decapitation, fixing the heads as described in the following section.

#### MK-801 Treatment

A pair of adult male zebrafish of similar size was placed in the experimental tank, separated by a barrier that isolated them visually but not chemically. After 24 hours of isolation, the barrier was removed, and their interaction was recorded for 30 minutes. After this time, they were separated and placed for 60 minutes in a treatment tank with MK-801 dissolved in water, as described in Cavallino et al. 2024. Then, they returned to their respective experimental tank, where they remained isolated for 48 hours until the barrier was removed, allowing a new interaction between the same opponents. After the 30-minute encounter, they were immediately sacrificed by cold shock and decapitation, fixing the heads as described in the following section.

### c-fos Immunohistochemistry

Immediately after the final encounter, individuals were euthanized by a rapid chilling followed by decapitation (American Veterinary Medical Association, AVMA, Guidelines for the Euthanasia of Animals, 2020). Histological processing of the brains was carried out following the protocol of Scaia et al., 2022. Briefly, heads were fixed in 10% formalin in PBS for three days. They were rinsed twice in PBS for 30 minutes each and decalcified in EDTA for two days. After two rinses in distilled water, samples were dehydrated in increasing concentrations of alcohol (70%, 96%, 100%), alcohol/xylene, xylene, and then two periods of 1:30 hours in paraffin I and II. Once embedded, they were mounted in paraffin blocks for histological processing. Coronal sections were cut at 7-micrometers using a microtome (Microm HM 350) and gelatin-coated slides. Sections were deparaffinized, hydrated, and rinsed before immunohistochemistry. The immunohistochemistry protocol began with blocking endogenous peroxidases (3% hydrogen peroxide for 5 minutes), rinsing, and then blocking nonspecific protein binding sites (5% skim milk for 1 hour at room temperature).

Slides were then rinsed and incubated at room temperature for 90 minutes with the primary antibody (polyclonal Goat Anti c-fos Abcam (Ab156802) 1/100 in PBS), followed by overnight incubation at 4°C. For the negative control, sections were incubated in the same volume of PBS instead of the primary antibody, following the same protocol. The following day, slides were removed from refrigeration and incubated for an additional 1 hour at room temperature. After PBS rinsing, sections were incubated for 1 hour at room temperature with the secondary biotinylated antibody (Rabbit antigoat DAKO (E0466) 1/200 in PBS). Sections were rinsed and incubated for 1 hour at room temperature in Streptavidin HRP (Chemicon (SA202) 1/400 in PBS), rinsed again and developed using DAB (3,3′-Diaminobenzidine, Cell marque 957D-20). The DAB reaction was terminated with distilled water, followed by counterstaining with hematoxylin. Samples were dehydrated and mounted in Canada balsam. After completing this protocol, photographs of the samples were taken with an Axiocam 208 color camera attached to a Zeiss Primo Star microscope for subsequent analysis and quantification (Supplementary material Fig.S1).

#### Image Analysis

Five coronal sections from each region were selected for analysis for each brain, leaving one section between them to avoid multiple quantifications of the same cells. Thus, five cross-sectional slices were quantified for each region per individual. The regions of interest, Vv, Vd, Dm, and Dl, were determined following the species atlas (Wulliman et al., 2012). Image analysis and c-fos positive cell quantification were carried out following a protocol established by Scaia et al. (2022). Within each region, two 1000 μm² rectangles were delineated for each slice, encompassing areas with the highest density of immunopositive cells. The number of immunopositive cells for c-fos within each rectangle was quantified, summing the values obtained for all sections, to finally get the number of c-fos cells in a 10,000 μm² area for each brain region. Quantification was performed in a single brain hemisphere, typically the one with the highest number of immunopositive cells. The total area of this hemisphere was quantified to normalize the number of c-fos immunopositive cells per total area of the brain hemisphere for each individual. Image analysis was conducted using Zen 3.3 Blue edition software.

#### Data Analysis

The number of c-fos immunopositive cells per analyzed region (relative to the area of the cerebral hemisphere) was compared between each treatment using a Non-parametric analysis, the Kruskal-Wallis test. In cases where Kruskall-Wallis was significant, a post hoc Dunn’s test was performed to compare among treatments. All median data refer to the number of c-fos-immunopositive cells in a 10,000 μm² region divided by the total area (TA) of each analyzed cerebral hemisphere section (c-fos IC/TA). The data are presented as median and interquartile range (IQR). Statistical significance was set at p < 0.05. A non-parametric Wilcoxon paired analysis was performed to compare neuronal activation in winners and losers, suggesting no significant differences in MK-treated fish in all four areas (Supplementary material Table. S1).

A distance-based permutation multivariate analysis of variance (PERMANOVA, Anderson, 2001a; Anderson and Walsh, 2013) and Principal Component Analysis (PCA) were used to explore individual variability in c-fos immunopositive cells. PERMANOVA test was performed on Euclidean distance matrices on all four brain areas as variables, with 999 random permutations (Anderson, 2001b).

A functional connectivity analysis was also conducted to evaluate the correlation between nuclei. Spearman correlation matrices were calculated for each treatment. Total connectivity (sum of all the correlations), Strength centrality (sum of all correlations for each nucleus), and Eigenvector centrality (taking into account the connections of each nucleus and the importance of the nuclei to which it is connected) were calculated. These correlations and their distributions were compared between treatments using the Kruskal-Wallis test followed by Dunn’s test, Mann-Whitney U (MWU) test, the Kolmogorov-Smirnov (KS) test, and the permutation test similar to the analysis performed by Scaia et al (2022).

Analyses were done in Python 3.11 and R 4.4.1. Boxplots were performed in GraphPad Prism 6. Scripts and data are available from the corresponding authors upon request.

## Results

As we described in a previous report, individuals treated with water after the first encounter resolved the second encounter with lower levels of aggression (a 67% decrease). At the same time, those treated with MK-801, an amnesic agent, did not significantly decrease their aggression (34%) (Cavallino et al., 2024). Interestingly, this maintenance of aggression levels was observed when the second encounter was against an unfamiliar opponent (Cavallino et al., 2020). In all cases, the winner of the first encounter also won the second.

Taking this into account, and as we suggested in our previous work, the individuals treated with water would recognize the opponent and/or the characteristics of the previous encounter and resolve a second encounter with lower aggressiveness. Individuals treated with an amnesic agent after the first encounter does not decrease their levels of aggressiveness in the second fight. Instead, they behave as if facing a novel opponent, showing they would not remember the opponent in particular. Here, and going a step forward, brain samples were processed for brain activation analysis to assess the neural correlate of these behaviors in Water and MK-801-treated fish.

### Ventral Nucleus of the Ventral Telencephalic Area (Vv)

When comparing the number of c-fos immunopositive cells per total area among the three treatments (Control, Water, and MK-801), significant differences were found (Kruskal-Wallis chi-squared = 9.357, df = 2, p-value = 0.0092). Individuals not exposed to social encounters in Control group show reduced immunopositive cells when compared to both groups of fish participating in social encounters under the Water treatment (Control: median: 0.000148 c-fos IC/TA, IQR: 0.000078, n=6; Water: median: 0.000264 c-fos IC/TA, IQR: 0.000028, n=6; p-value = 0.0060), as well as in the MK-801 treatment MK-801 (MK-801: median: 0.000250 c-fos IC/TA, IQR: 0.000031, n=9; p-value = 0.0084). No differences were found between the Water and MK-801 treatments (p-value = 0.7085) Fig. 2A.

**Figure 1.**
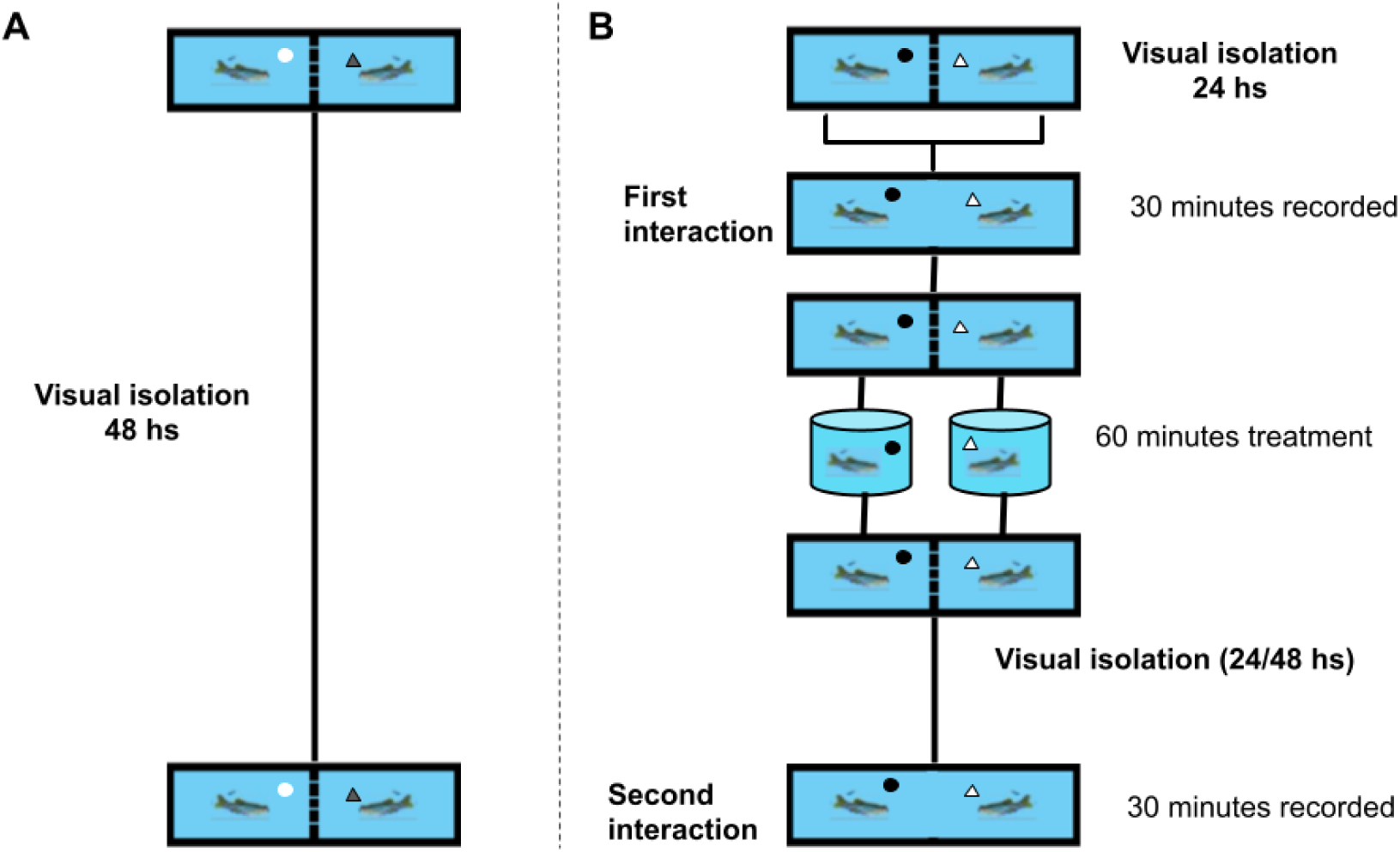
Experimental device and treatments. A) pairs of zebrafish males were isolated in a 2 L aquarium, separated by a barrier. Forty-eight hours later, individuals were euthanized by a cold shock and decapitation. B) Experimental devices in which pairs of zebrafish males were isolated in a 2 L aquarium, separated by a barrier. Twenty-four hours later, individuals were allowed to interact for 30 minutes until separated again. Immediately after the first encounter, pairs were moved to separate aquaria for 60 minutes, containing water or MK-801 dissolved in water. After treatment, pairs were returned to their tank and isolated for 24 or 48 hours until they interacted again for 30 minutes. After this encounter, individuals were euthanized by a cold shock and decapitation.

**Figure 2.**
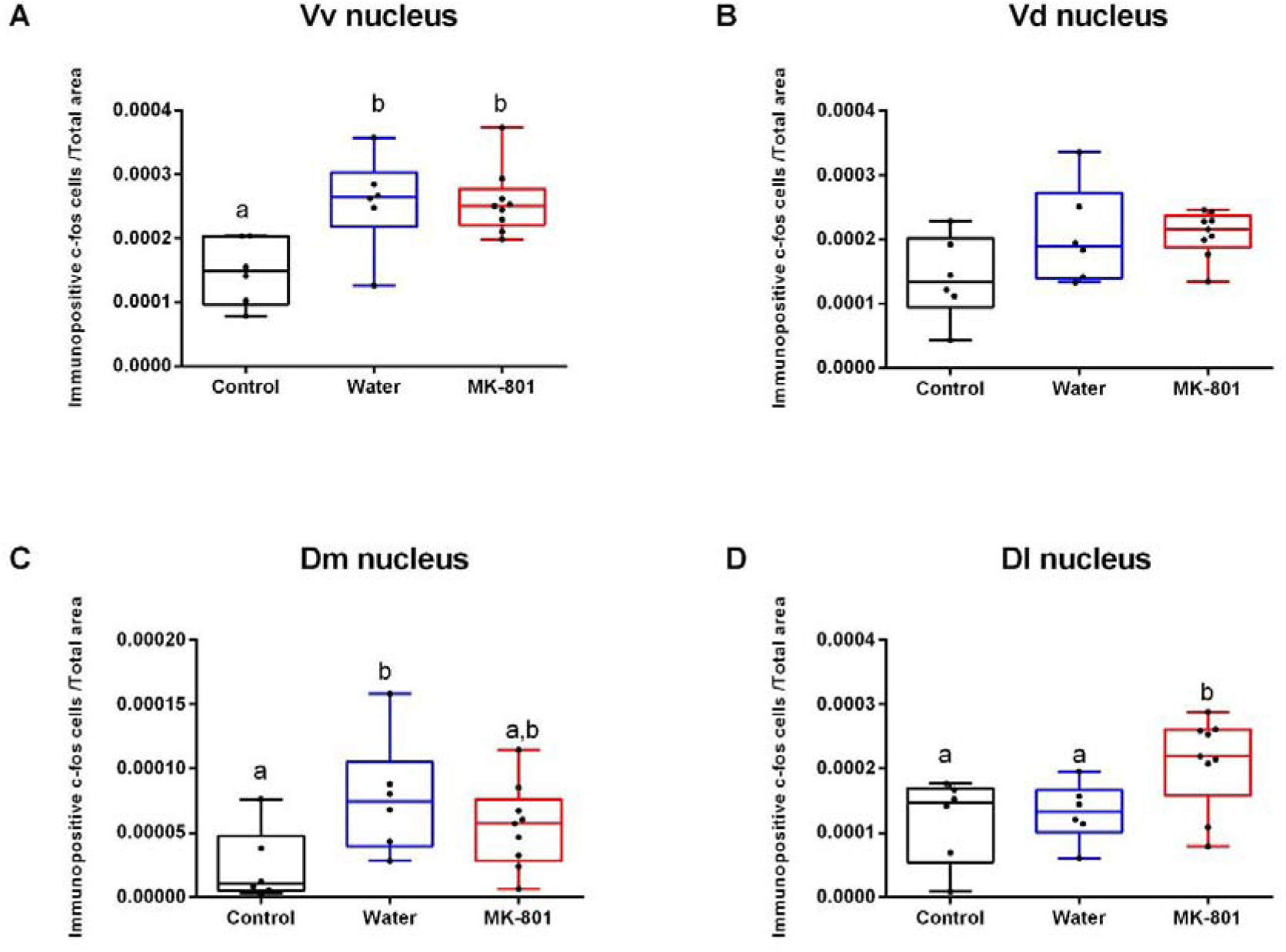
Neuronal activation of each treatment for the analyzed nuclei. Boxplot for the number of c-fos immunopositive cells/ total area of the cerebral hemisphere (IC/TA) for the A) Vv nucleus. B) Vd nucleus. C) Dm nucleus and D) Dl nucleus comparing the Control, Water, and MK-801 treatments. The band inside the box represents the median, and the whiskers represent the maximums and minimums. The circles represent the value of each individual. Different letters indicate significant differences between treatments.

### Dorsal Nucleus of the Ventral Telencephalic Area (Vd)

When comparing the number of c-fos immunopositive cells per total area among the three treatments (Control, Water and MK-801), no significant differences were found (Kruskal-Wallis chi-squared = 4.6608, df = 2, p-value = 0.0972) (Median: Control 0.000132 c-fos IC/TA, IQR: 0.000066; Water 0.000188 c-fos IC/TA, IQR: 0.000085; MK-801 0.000215 c-fos IC/TA, IQR: 0.000030) Fig. 2B.

### Medial Nucleus of the Dorsal Telencephalic Area (Dm)

When comparing the number of c-fos immunopositive cells per total area among the three treatments (Control, Water, and MK-801), significant differences were found (Kruskal-Wallis chi-squared = 6.795, df = 2, p-value = 0.0334). Individuals not exposed to social encounters in the Control group show reduced immunopositive cells when compared to fish participating in social encounters and exposed to water (Control: median: 0.000012 c-fos IC/TA, IQR: 0.000025, n=6; Water: median: 0.000074 c-fos IC/TA, IQR: 0.000036, n=6; p-value = 0.0105). Although no significant differences were found between Control and MK-801 treatment, there is a trend with a p-value close to the defined significance threshold (MK-801: median: 0.000057 c-fos IC/TA, IQR: 0.000023, n=9; p-value = 0.0691). No differences were found between the Water and MK-801 treatments (p-value = 0.3244) Fig. 2C.

### Lateral Nucleus of the Dorsal Telencephalic Area (Dl)

When comparing the number of c-fos immunopositive cells per total area among the three treatments (Control, Water, and MK-801), significant differences were found (Kruskal-Wallis chi-squared = 6.5541; df = 2, p-value = 0.0377). Individuals not exposed to social encounters in the Control group show no differences in neural activation when compared to fish participating in social encounters and exposed to water (Control: median: 0.000146 c-fos IC/TA, IQR: 0.000076, n=6; Water: median: 0.000132 c-fos IC/TA, IQR: 0.000037, n=6; p-value=0.9285), they show reduced immunopositive cells when compared to the MK-801 group (MK-801: median: 0.000219 c-fos IC/TA, IQR: 0.000045, n=9; p-value = 0.0284). Fish exposed to water showed reduced c-fos immunopositive cells in Dl compared to those exposed to MK-801 (p-value = 0.0366) Fig. 2D.

In sum, the results revealed significant differences in neuronal activation across various brain nuclei based on the specific treatment administered. In particular, there was greater activation of the Vv nucleus in fish exposed to social encounters and a significant difference in the activation of the Dl nucleus depending on whether they had been treated with water or the amnesic agent. Nevertheless, this is an initial attempt to find differences; more evidence is needed to corroborate the differences between these treatments. Therefore, we performed the following analysis.

### Individual patterns of neuronal activation across integrated brain nuclei

In order to explore individual variability in neuronal activation and to assess whether data can be clustered in different groups corresponding to treatments, PCA followed by Permanova was performed. The first and second components explained a total variance of 84.14% (PC1 67.65% and PC2 16.5%). Table 1 indicates component interpretations, showing that areas corresponding to the ventral telencephalon (Vd and Vv) show higher loadings for PC1, whereas areas from the dorsal telencephalon (Dm and Dl) contribute more to PC2 (Fig. 3, Table 1). We found significant differences in PC loadings between treatments when a perMANOVA was performed (F=5.3926, P-value=0.006, 999 permutations). The Control group differed from MK-801 and Water treatment (Control vs. MK-801 p=0.024; Control vs. Water p=0.039); no significant differences were found between MK-801 and Water (p=0.624).

**Figure 3.**
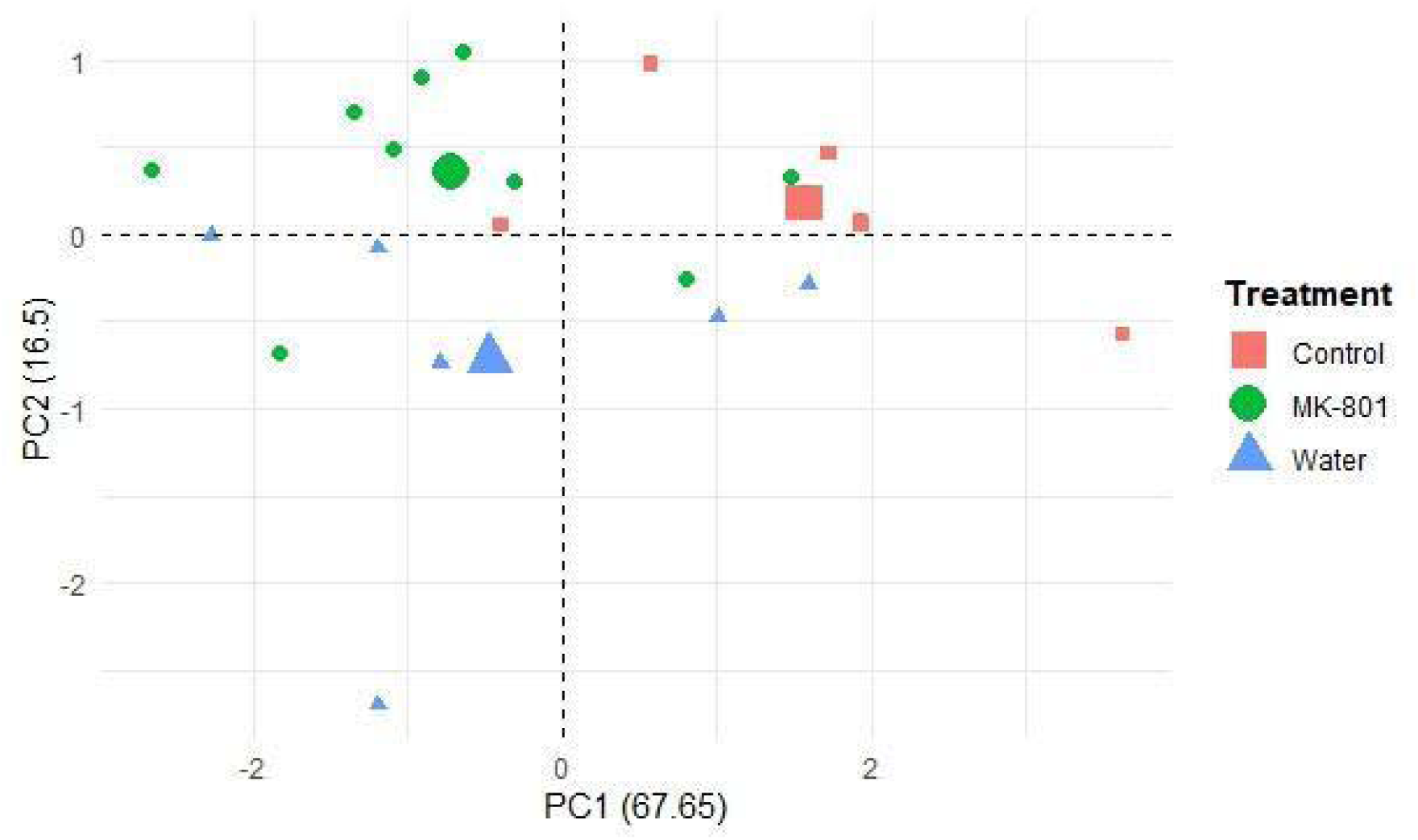
Principal component analysis for the Control, MK-801, and Water treatment. Red squares indicate individuals’ scores for the Control group; green circles indicate individuals’ scores for MK-801 treatment, and blue triangle indicates individuals’ scores for Water treatment. The largest symbols represent the centroid of each treatment.

**Table 1.** The proportion of variance indicates the percentage of variance of the nucleus activation explained by each principal component (PC). The cumulative proportion shows the explained variance of the PC plus the preceding PC. For each nucleus (Vv, Vd, Dm and Dl), the loadings are presented. The higher loadings for each component are shown in bold.

|  | PC1 | PC2 |
| --- | --- | --- |
| Standard deviation | 1.6449 | 0.8123 |
| Proportion of variance | 0.6765 | 0.1650 |
| Cumulative proportion | 0.6765 | 0.8414 |
| Vv | **-0.5392** | -0.0615 |
| Vd | **-0.5551** | 0.2240 |
| Dm | -0.4271 | **-0.8073** |
| DI | -0.4675 | **0.5424** |

This analysis considering all four brain areas as variables suggests differences in neuronal activation between individuals exposed to the physical interaction (Water and MK-801 treatments) and those not exposed to an opponent, which can be separated and visualized in a principal component analysis. The next step was to evaluate whether functional connectivity among these nuclei differs in the experimental groups.

### Functional connectivity: Neuronal correlation

To evaluate not only the individual activation of each brain nucleus but also the relationships among them, functional connectivity networks were constructed for each experimental condition based on co-activation matrices on the brain regions analyzed. For this, Spearman correlations for the number of c-fos immunopositive cells (c-fos IC/TA) among all the analyzed nuclei (Vv, Vd, Dm, Dl) were calculated for each treatment (Fig. 4).

**Figure 4.**
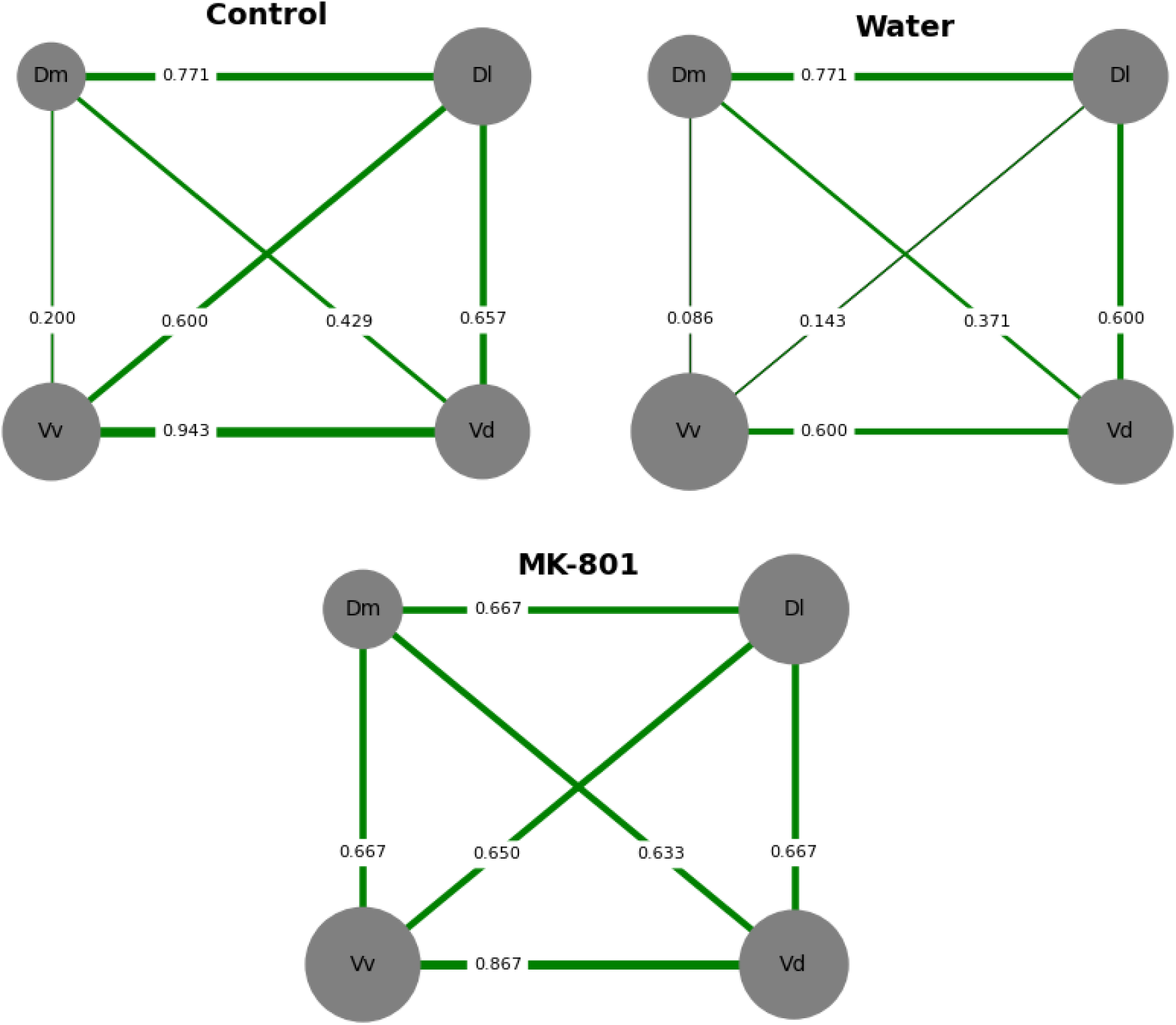
Neuronal correlation of the four nuclei (Vv, Vd, Dm, and Dl) for the three treatments (Control, Water, and MK-801). All the Spearman correlations were positive. The size of each nucleus was proportional to their activation (c-fos IC/TA), and the thickness of the connection lines was proportional to the level of interaction between each pair of nuclei (Spearman’s correlation coefficient). Numbers in each line indicate the Spearman’s correlation coefficient.

The Spearman correlation matrices for each treatment are shown in Supplementary Material Table S2. Results suggest all positive correlations (i.e., functional co-activation between brain regions), with no negative correlations for any pair of nuclei (i.e., functional co-inhibition between brain regions).

Significant differences were found when comparing the MK-801 and Water treatments for both the KS test and the MWU test (KS test: statistic = 0.833, p-value = 0.025; MWU test: statistic = 5.0, p-value = 0.0434). No significant differences were found between the Control and Water treatments (KS test: statistic = 0.333, p-value = 0.930; MWU test: statistic = 10.5, p-value = 0.258) or between the Control and MK-801 treatments (KS test: statistic = 0.5, p-value = 0.474; MWU test: statistic = 13.0, p-value = 0.468). Similar results were found when Dunn’s test was performed (MK-801 vs. Water: P-value=0.0364; MK-801 vs. Control: P-value=0.3997; Water vs. Control: P-value=0.2114). Nevertheless, no differences were found when the permutation test was performed (MK-801 vs. Water: P-value=0.250; MK-801 vs. Control: P-value=0.762; Water vs. Control: P-value=0.368).

### Functional connectivity: Total connectivity, strength, and eigenvector centrality

When analyzing total connectivity, although there appears to be a trend showing higher connectivity for the MK-801 treatment (Total connectivity: 4.15) compared to Water (Total connectivity: 2.57) and Control (Total connectivity: 3.6), no significant differences were found for any treatment (Supplementary material Table S3).

However, when analyzing strength centrality, a significant difference was found between treatments (Kruskal-Wallis chi-squared = 6.038; df = 2, p-value = 0.0488, followed by a Dunn’s test). Nuclei in MK-801 exhibited higher strength centrality than those in the Water treatment (p-value=0.0142). There were no differences between MK-801 and Control (p-value = 0.2807) or between Water and Control (p-value = 0.1698). Similar results were found when the permutation test was performed (MK-801 vs. Water: P-value=0.010; MK-801 vs. Control: P-value=0.422; Water vs. Control: P-value=0.084).

Each nucleus’s strength and eigenvector centrality were calculated and ranked according to their importance from highest to lowest scores. The three treatments differed in ranking the importance of their nuclei for both strength and eigenvector centrality. When analyzing the first position, the Vd nucleus was most relevant for the Water and Control treatments, and the Vv nucleus was most relevant for the MK-801 treatment. However, when considering the two most central nuclei in each experimental group, results suggest that while Vd is central for all experimental groups, Dl is highly relevant for Control and Water treatments and Vv is only relevant for the MK-801 treatment. Additionally, the Dm nucleus was the least relevant for the MK-801 and Control treatments, while the Vv nucleus was the least relevant for the Water treatment. The ranking is shown in Supplementary Material Table S4.

In this regard, connectivity throughout different brain nuclei seems to differ among treatments. In particular, results suggest significant differences between the Water and MK-801 treatments when comparing correlation matrices and a non-significant trend in overall connectivity. Additionally, results suggest a mild difference in strength centrality between experimental groups. Furthermore, even if comparison of centrality of each brain area was not statistically assessed, the order of importance of brain nuclei seems to differ in all three treatments, considering both strength and eigenvector centrality.

## Discussion

The main goal of our work was to evaluate the neuronal activity that underlies recognition of an opponent and long-term memory associated with social interactions. To disentangle which is the neural substrate of these processes throughout the SDMN we analyzed the neuronal activation of different brain nuclei associated with aggression, memory, and social interactions in a social memory paradigm, by quantifying the c-fos expression in male zebrafish. We found a higher activation in the Dl nucleus in MK-801-treated fish, and a lower activation in the Vv nucleus in the Control group. Furthermore, we were able to differentiate between individuals that had physical interaction (Water and MK-801 treatment) and those that did not (control group) through a PCA analysis and PERMANOVA.

It is important to note that the individuals were sacrificed immediately after the 30 minutes following the start of the interaction (removal of the barrier) of the second encounter in the Water and MK-801 treatments, while the Control group was sacrificed after undergoing similar manipulations but without being exposed to the agonistic encounter. Considering the time required for c-fos expression (Chaudhuri et al., 2000), this method does not quantify activation corresponding to the agonistic encounter and resolution on the second encounter. In this way, the effect we are detecting on neuronal activation is the result of this initial interaction (or none in the case of the Control), meaning that we are detecting changes associated to the first interactions with an individual who can be recognized and remembered, depending on whether a memory of the previous encounter was formed. In the case of an amnesic effect (MK-801 Treatment), and considering the already reported effects on behavioral outcome (Cavallino et al., 2024), neuronal activation observed in this group does not account for recalling the first encounter, but for a novel interaction between two unfamiliar individuals instead.

As we mentioned before, c-fos protein levels can be observed between 30 and 120 minutes after the stimulus, with a maximum peak at 90 minutes (Kovács, 1998; Chaudhuri et al., 2000; Von Trotha et al., 2014; Chatterjee et al., 2015). In the present work, by sacrificing the individuals immediately after the 30-minute interaction, we would not be observing the effect of fight resolution and aggressive displays *per se* but rather the impact of the initial moments of interaction. During these moments, specific nuclei related to social interaction and those associated with recalling the memory formed by the previous encounter (individuals with prior experience against that opponent) could be activated. However, if the stimulus is perceived as novel (e.g., in individuals treated with MK-801 after the first encounter), we could observe activation related to the first steps of the evaluation and recognition of that particular individual. For this reason, we decided to analyze protein expression 30 minutes after the first interaction of the second encounter, thereby avoiding the confusion of effects with activation due to the fight and aggressive displays themselves. Nonetheless, it would be interesting to evaluate c-fos expression at longer time points, such as 60 minutes, encompassing the peak of expression, and 90 minutes, where we would observe the effect of the agonistic encounter and resolution, to compare them with results reported here.

We aim to evaluate brain activity in specific nuclei described to be associated with aggression, emotional learning, and memory. Teles and colleagues (2015) found that, when comparing c-fos mRNA expression for the Dm, Dl, Vv, Vs, and preoptic area (POA) nuclei, zebrafish visually isolated from a conspecific showed lower expression levels in all these nuclei compared to individuals exposed to an agonistic encounter or a mirror interaction (Teles et al., 2015). These areas are part of the SDMN and represent promising regions to evaluate activation concerning agonistic encounters. In that study, the authors report the effect of an agonistic interaction and all the aggressive behaviors involved in brain activation. In our work, we evaluate the activation related to the first moments of interaction in Vv, Vd, Dm, and Dl nuclei.

For the ventral region of the telencephalon, we found that in the ventral nucleus (Vv), individuals who were not exposed to a social agonistic encounter showed fewer c-fos immunopositive cells than those who did have this interaction, regardless of whether they were treated after the first encounter with MK-801 or water. Although no significant differences were found for the dorsal nucleus of the ventral area (Vd), there seemed to be a trend towards the same result. This evidence suggests that these nuclei, particularly Vv, might be involved in social interactions, regardless of whether it is with a familiar individual. This interpretation arises because, even if behavioral results suggest that fish treated with an amnesic agent did not recognize their opponent but fish treated with water were able to recognize them (Cavallino et al., 2024), the same levels of neuronal activation in Vv and Vd were observed in both groups. Regarding the Vv nucleus, Fan and colleagues evaluated neuronal activation related to social learning in guppies (*Poecillia reticulata*) by quantifying Ps6 (Fan et al., 2022), a ribosomal protein in which phosphorylation indicates neuronal activation (Knight et al., 2012). Therefore, although this is not an IEG, this protein has been widely used to evaluate neuronal activation in various ethological contexts to study different aspects of social behavior in fish (Akinrinade et al., 2023; Scaia et al., 2022; Nunes et al., 2021). Fan and colleagues found increased Ps6 immunolabeling in the POA and the Vv nucleus during the acquisition of social learning related to a threat (via an alarm chemical cue from conspecifics), highlighting the importance of the Vv nucleus in this type of learning (Fan et al., 2022). Similar results were found in zebrafish; Pinho and colleagues (2023) found that in a classical conditioning paradigm, an increase in activity in Vv (through quantification of c-fos expression) was related to social learning but not with asocial learning. Our findings further emphasize the significant involvement of the Vv nucleus in social learning, which corroborates previous evidence supporting this conclusion.

Regarding the brain areas in the dorsal telencephalon , the medial nucleus (Dm) showed similar results than the Vv with lower activation in the Control group, but in this case no differences were found between fish previously treated with MK-801 and those treated with water. However, in the lateral nucleus (Dl) we found more c-fos immunopositive cells in animals treated with MK-801 the previous day when compared to those exposed to water or controls, with no differences between these two. These results suggest that the Dl might be involved in the novel recognition of an individual but not in repeated exposure to the same opponent.

These differences can be better understood considering that while in this study we quantified c-fos protein, Teles et al assessed c-fos mRNA which does not always correlate with protein levels due to post-transcriptional modifications (Chatterjee et al., 2015). Moreover, while Teles and colleagues observed c-fos levels related to the effects of fighting between individuals, this study refers to c-fos expression due to the recognition of a previous opponent during the first moments of a second interaction, as already discussed. Finally, there might be an effect related to the repetitiveness of stimuli, as we will discuss below. These findings on neuronal activation, together with our previous behavioral results (Cavallino et al., 2024), suggest that activation of Dl nucleus can be triggered in response to a novel stimulus (such as the presentation of an unfamiliar individual) and could be involved in the formation of long-term memory related to this novel individual, however, it might not respond in the same way during memory recall. Nevertheless, it would be necessary to increase the amount of data to enhance the robustness of the analysis. It is important to note that the activation and response we are referring to is through c-fos expression, as observed in other studies where not all immediate-early genes respond the same to the same stimulus. Teles and colleagues found differences depending on whether they evaluated c-fos or other immediate-early genes, egr1 (Teles et al., 2015).

In line with our results, studies in other species report differential activation on whether a stimulus is presented for the first time or as a result of several presentations. For example in the common sparrow, Kimball and colleagues observed that exposure to a series of objects during several days increased c-fos immunolabeling in the caudal hippocampus and the ventral medial arcopallium (homologous to some areas of the mammalian amygdala) only when presented with a novel object the next day, one they had not been exposed to before. However, this increase was not observed if presented with an object they had already been exposed to. This suggests that these regions might be involved in recognizing novel objects but would not activate through c-fos with repeated exposure to an object (Kimball et al., 2022). Similar evidence was found in male albino rats, where individuals without exposure to a maze showed low levels of c-fos expression and another IEG, C-jun, through immunohistochemical techniques. These expression levels increased considerably when the rats were exposed to a maze for the first time, but the increase was minor with repeated exposure. This effect was observed in various brain regions, including the hippocampus, somatosensory cortex, and cerebellum (Papa et al., 1993).

It is interesting to note that some studies in fish suggest that the Dl nucleus is mainly associated with spatial learning. In goldfish, active avoidance experiments using conditioning between a green light and an electric shock suggest that lesions in the Dm, but not in the Dl, induced a deficit in testing this conditioning. Conversely, authors observed the opposite in spatial learning experiments: lesions in the Dm did not affect conditioning, but in the Dl did. These results suggest that Dl is involved in spatial learning, while Dm is involved in emotional learning (Portovella et al., 2002). Overall, results in our work suggest that the Dl nucleus could be involved in social learning, particularly in recognizing a novel individual. However, it will be interesting to confirm the role of Dl in recognizing a novel opponent with other experimental approaches such as lesions and different ethological contexts.

When considering all the nuclei together, we found that individual variability in neuronal activation can be clustered according to different treatments. In particular, we could distinguish between individuals not exposed to physical interaction (Control group) and those that were involved in social interactions, regardless of the treatment (MK-801 or Water). These results show the effect of physical interactions on the telencephalic nuclei’s overall activity. Nevertheless, functional connectivity analysis suggests differences between individuals exposed to physical interaction according to the treatment. Individuals exposed to a non-remembered opponent showed higher strength centrality and a trend in higher connectivity among their nuclei and total connectivity when compared to individuals exposed to remembered opponents. These results suggest that different telencephalic nuclei and neural networks may be differentially activated depending on whether fish are exposed to a known conspecific they do not remember, or to an opponent that is already known and also remembered. It is important to point out that all the correlations found were positive, which could indicate a co-activation between all the nuclei. However, negative correlations could be found when evaluating other nuclei belonging to the SDMN, as seen in other studies in zebrafish (Scaia et al., 2022).

Considering that the Dm nucleus is the suggested homology of the basolateral amygdala of mammals, which is involved in aggression, emotional learning, and memory (O’Connell and Hofmann, 2011; Von Trotha et al., 2014), and previous results in fish suggesting that this region would be involved in these functions (Martín et al., 2011; Lal et al., 2018; Lau et al., 2011), we hypothesized that there would be differences in the activation of this nucleus when comparing individuals exposed to a recognized and remembered opponent with those exposed to an unrecognized opponent. However, we did not find these differences, nor did it appear to be a prominent nucleus in terms of connectivity with the other analyzed nuclei.

Nevertheless, we do not rule out the possibility that assessing a greater number of nuclei belonging to the SDMN could reveal a more significant role for the Dm. Further experiments would be necessary to verify the influence of the nuclei analyzed here on an opponent’s recognition and memory formation, such as analyzing brain activation in individuals exposed to a novel opponent on the second day and comparing it with activation in the MK-801 treatment. Additionally, it would be interesting to analyze more nuclei and hypothalamic areas to observe a more enriched functional connectivity network.

Our results provide a first step in studying neuronal activation associated with the retrieval of a memory of a previous opponent and the effect of a first interaction with a non-remembered one. We postulate in a promising way that the Vv and Vd nuclei are associated with social interactions and that the Dl nucleus is relevant for individual recognition, as well as the importance of analyzing the connectivity between nuclei to be able to more fully evaluate the neural activity underlying social memory and recognition of a conspecific.

## Acknowledgement

To Matias Pandolfi for his contribution to this work, for starting this project and for all these years.

## Data and resource availability

Data that support these results are available from the corresponding authors upon request.

## Supplementary material

**Figure S1.**
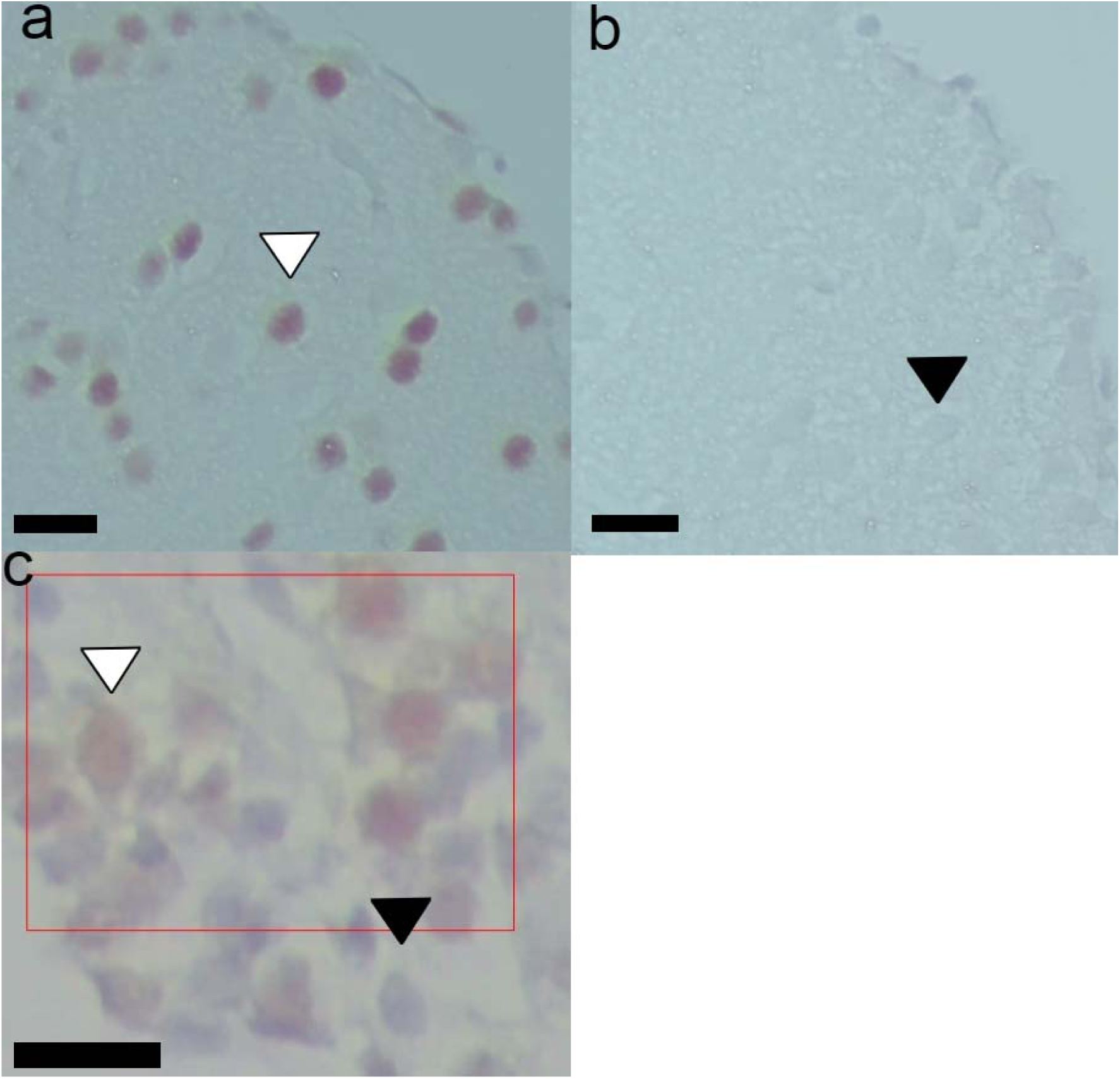
A portion of the Dl region is shown for A) positive label for c-fos and B) Negative control. Black triangles indicate unlabeled nuclei, and White triangles indicate positively labeled nuclei. 400 x magnifications. The black bar corresponds to 10 μm. C) The red rectangle represents one of the 1000 um2 regions in which the amount of c-fos immunopositive cells are quantified (White triangle); unlabeled nuclei can also be observed (Black triangle). 400 x magnifications. The black bar corresponds to 10 μm.

**Table S1.**
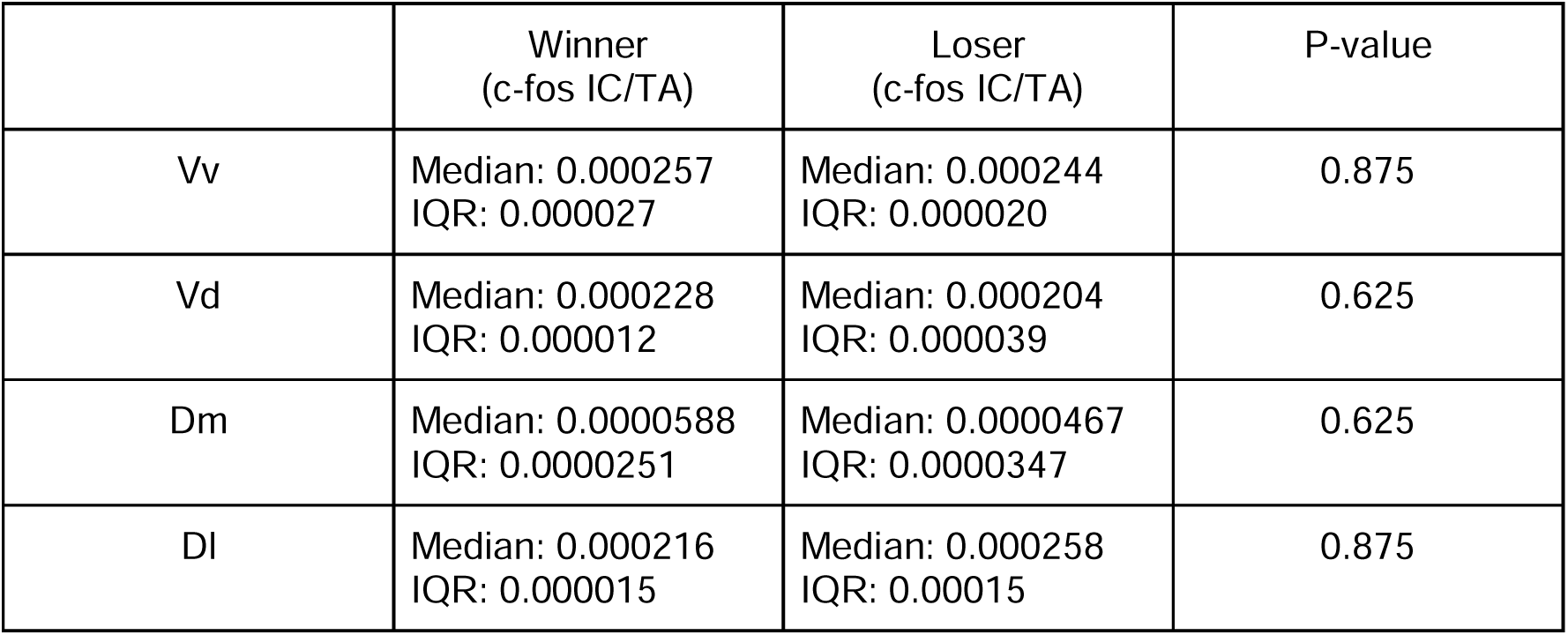
c-fos IC/TA for winners (n=4) and losers (n=5) for Vv, Vd, Dm, and Dl nucleus (MK-801 treatment). P-values for the Mann-Whitney test comparing winners and losers for each nucleus were shown.

**Table S2.**
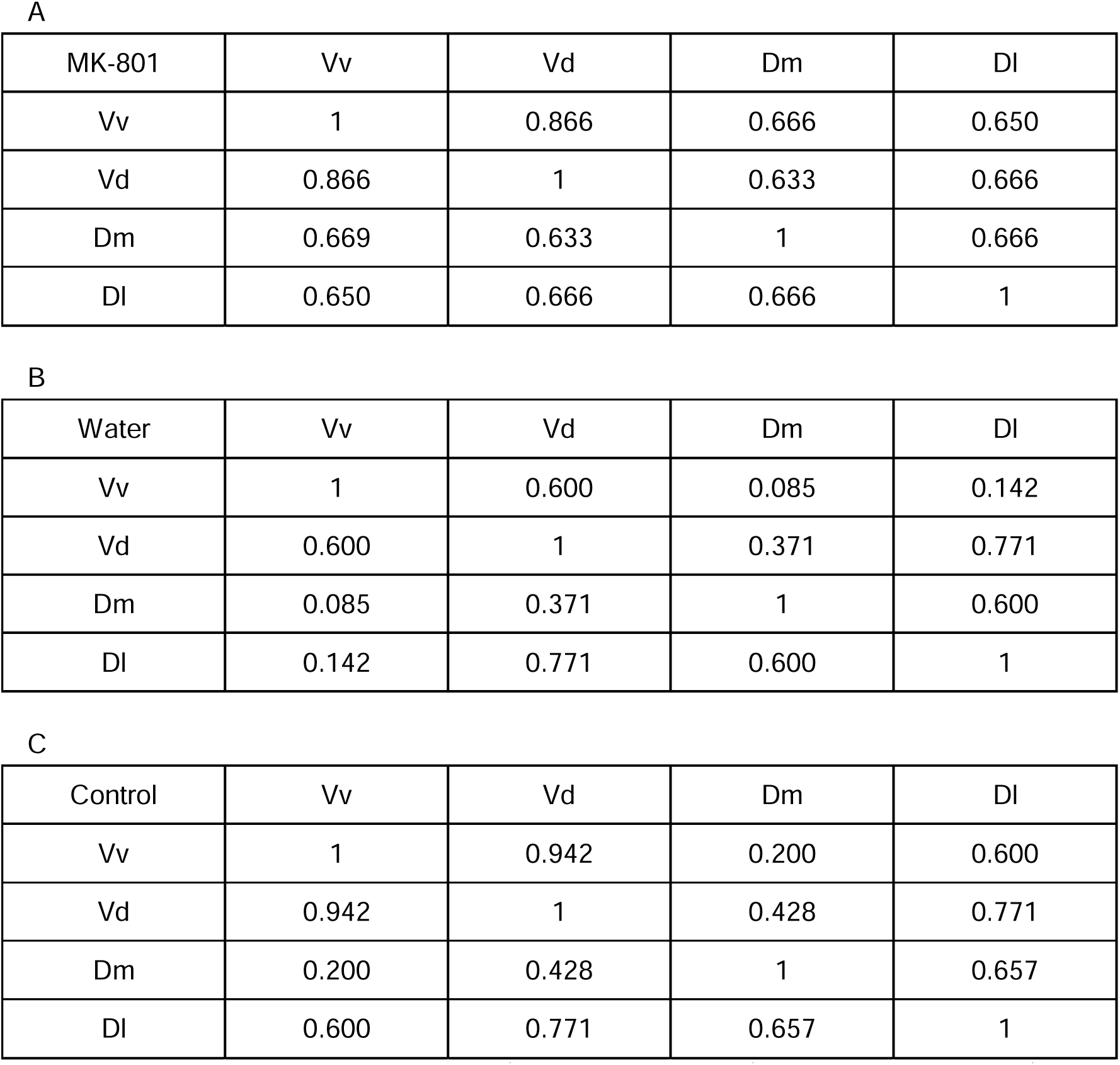
Spearman correlation matrix for A) MK-801 treatment, B) Water Treatment, and C) Control. Spearman correlation coefficient was shown for each pair of nuclei.

| MK-801 | Vv | Vd | Dm | DI |
| --- | --- | --- | --- | --- |
| Vv | 1 | 0.866 | 0.666 | 0.650 |
| Vd | 0.866 | 1 | 0.633 | 0.666 |
| Dm | 0.669 | 0.633 | 1 | 0.666 |
| DI | 0.650 | 0.666 | 0.666 | 1 |

B
| Water | Vv | Vd | Dm | DI |
| --- | --- | --- | --- | --- |
| Vv | 1 | 0.600 | 0.085 | 0.142 |
| Vd | 0.600 | 1 | 0.371 | 0.771 |
| Dm | 0.085 | 0.371 | 1 | 0.600 |
| DI | 0.142 | 0.771 | 0.600 | 1 |

| Control | Vv | Vd | Dm | DI |
| --- | --- | --- | --- | --- |
| Vv | 1 | 0.942 | 0.200 | 0.600 |
| Vd | 0.942 | 1 | 0.428 | 0.771 |
| Dm | 0.200 | 0.428 | 1 | 0.657 |
| DI | 0.600 | 0.771 | 0.657 | 1 |

**Table S3.**
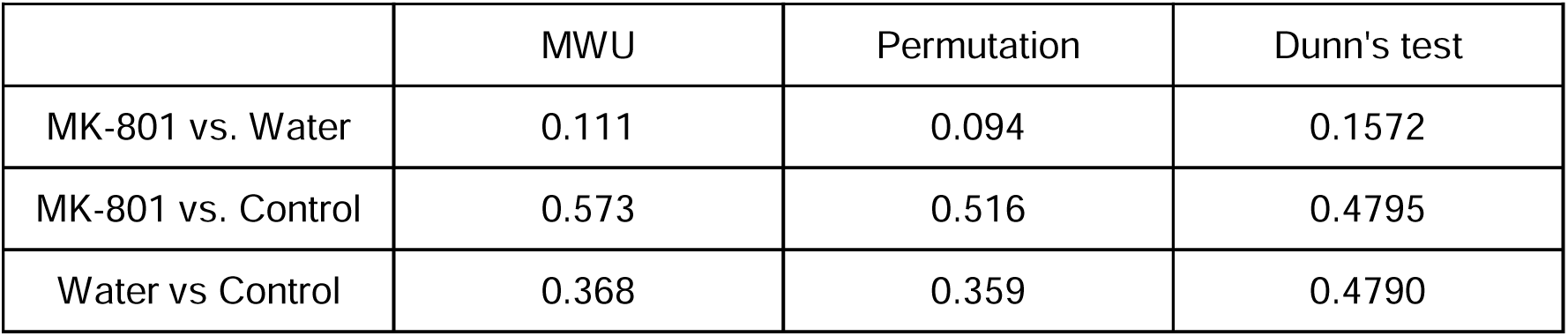
P-values of Mann-Whitney U, permutation, and Dunn’s test comparing total connectivity between each pair of treatments.

**Table S4.**
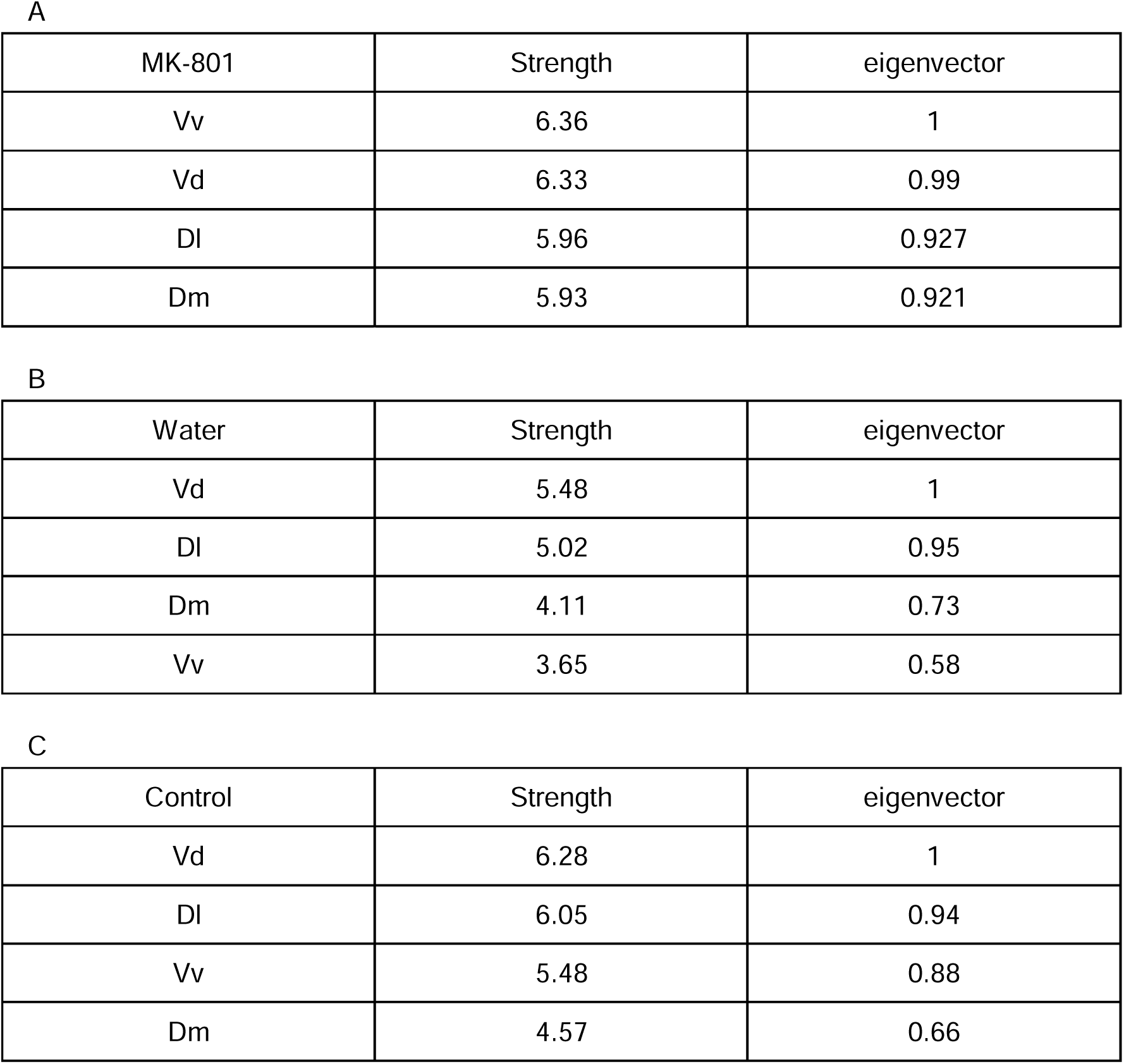
Strength centrality and eigenvector centrality for A) MK-801 treatment. B) Water treatment and c) Control. The nuclei were ordered from highest to lowest for each treatment.

| MK-801 | Strength | eigenvector |
| --- | --- | --- |
| Vv | 6.36 | 1 |
| Vd | 6.33 | 0.99 |
| DI | 5.96 | 0.927 |
| Dm | 5.93 | 0.921 |

B
| Water | Strength | eigenvector |
| --- | --- | --- |
| Vd | 5.48 | 1 |
| DI | 5.02 | 0.95 |
| Dm | 4.11 | 0.73 |
| Vv | 3.65 | 0.58 |

| Control | Strength | eigenvector |
| --- | --- | --- |
| Vd | 6.28 | 1 |
| DI | 6.05 | 0.94 |
| Vv | 5.48 | 0.88 |
| Dm | 4.57 | 0.66 |

